# Sterol O-acyltransferase (SOAT/ACAT) activity is required to form cholesterol crystals in hepatocyte lipid droplets

**DOI:** 10.1101/2024.01.20.576345

**Authors:** Jordan A Bairos, Uche Njoku, Maria Zafar, May G Akl, Lei Li, Gunes Parlakgul, Ana Paula Arruda, Scott B Widenmaier

**Author notes:** **Correspondence to Lead Contact:** 107 Wiggins Rd, Health Sciences Building Saskatoon, SK, S7N 5E5, Canada. Co-first authors (Jordan A Bairos, Uche Njoku).

## Abstract

**Objective:** Excess unesterified (free) cholesterol can induce formation of cholesterol crystals in hepatocyte lipid droplets. Presence of such crystal distinguishes metabolic dysfunction associated steatohepatitis (MASH) from simple steatosis and may underlie its pathogenesis by causing cell damage that triggers liver inflammation. The mechanism linking cholesterol excess to its crystallization in lipid droplets is unclear. As cholesteryl esters localize to and accumulate in lipid droplets much more readily than free cholesterol, we investigated whether cholesterol esterification by sterol O-acyltransferase (SOAT), also known as acyl co-A cholesterol acyltransferase (ACAT) is required for hepatocyte lipid droplet crystal formation.

**Method:** Cholesterol crystals were measured in cholesterol loaded Hep3B hepatocytes, RAW264.7 macrophages and mouse liver using polarizing light microscopy. We examined the effect of blocking SOAT activity on crystal formation and compared these results to cholesterol metabolism and the progression to intracellular crystal deposits.

**Results:** Cholesterol loading of Hep3B cells caused robust levels of lipid droplet localized crystal formation in a dose- and time-dependent manner. Co-treatment with SOAT inhibitors and genetic ablation of *SOAT1* blocked crystal formation. SOAT inhibitor also blocked crystal formation in low density lipoprotein (LDL) treated Hep3B cells, acetylated LDL treated RAW 264.7 macrophages, and in the liver of mice genetically predisposed to hepatic cholesterol overload and in mice fed a cholesterol enriched, MASH-promoting diet for 24 weeks.

**Conclusion:** SOAT1-mediated esterification may underlie cholesterol crystals associated with MASH by concentrating it in lipid droplets. These findings imply that inhibiting hepatocyte SOAT1 may alleviate cholesterol associated MASH. Moreover, that a lipid droplet localized cholesteryl ester hydrolase may be required for cholesterol crystal formation or, instead, that the crystals are composed of cholesteryl ester.

**Funding Sources:** Grants supporting this research were awarded to SBW from the Natural Sciences and Engineering Research Council of Canada (NSERC). SBW was supported by a National New Investigator Award and McDonald Scholarship from the Heart and Stroke Foundation of Canada. UN and MA were supported by a James Regan Cardiology Research scholarship from University of Saskatchewan’s College of Medicine.

## 1. INTRODUCTION

Cholesterol is essential for cell membrane homeostasis [1]. However, excess unesterified (free) cholesterol can exert damaging and stressful effects on cellular processes that promote disease [2]. One disease linked to excess free cholesterol is metabolic dysfunction-associated steatohepatitis (MASH), also known as non-alcoholic steatohepatitis [3, 4]. Free cholesterol in liver is elevated in people with MASH compared to simple steatosis [5–7] and rodent studies show hepatic cholesterol accumulation triggers the progression of steatosis to MASH [4, 8–14]. Deciphering the link between hepatic cholesterol excess and MASH may provide insight regarding its pathogenesis and reveal a strategy for uncoupling this link to improve liver health outcomes for at risk patients.

The mechanism by which excess hepatic cholesterol underlies MASH may involve the alteration of membrane composition and fluidity, induction of oxidative damage, accumulation of toxic oxysterols, and cholesterol-induced taffazin signaling [1–3, 15]. It is also known that free cholesterol exceeding the cell membrane carrying capacity [16] is prone to precipitate and form crystals, and that such crystals are linked to MASH [3]. Indeed, Ioannou and colleagues have shown excess cholesterol promotes the formation of lipid droplet localized cholesterol crystals in hepatocytes and that these crystals distinguish MASH from simple steatosis in humans and rodents [8, 10, 14]. Although the impact of such crystals on hepatocytes has not been established, atherosclerosis studies indicate that subintimal plaque-associated cholesterol crystals cause membrane damage and activate the inflammasome to promote atherothrombosis [17, 18], and it has been proposed that hepatocyte cholesterol crystals promote MASH in a similar manner [3].

Although free cholesterol excess and lipid droplet localized cholesterol crystals have independently been linked to MASH, the mechanism by which cholesterol accumulation results in the formation of lipid droplet localized cholesterol crystals is unclear. One proposal is that free cholesterol traffics to lipid droplets via membrane sites that are in contact with cholesterol overloaded organelles such as the endoplasmic reticulum (ER), late endosomes, and lysosomes [3, 14]. Accordingly, since lipid droplet membranes are a monolayer rather than bilayer there is a relatively low cholesterol carrying capacity, which may predispose cholesterol to precipitate and form crystals. One paradox however is that it is much more energetically favorable for free cholesterol to accumulate in the plasma membrane or other bilayer membranes, as they have higher carrying capacity [16]. Thus, free cholesterol flow to lipid droplets is likely miniscule, raising the possibility that it may concentrate at the lipid droplet via a different mechanism.

As hepatocytes are equipped with constraints that support membrane homeostasis as well as with adaptive stress averting responses that are engaged when free cholesterol exceeds a homeostatic threshold [1, 2, 16, 19], we considered whether these responses may contribute to the formation of cholesterol crystals. Here, we focus on the role of sterol O-acyltransferase (SOAT) activity. SOAT1 and SOAT2 are ER-localized enzymes that produce cholesteryl ester using free cholesterol and long chain fatty acyl coenzyme A as substrates [20, 21]. In hepatocytes, depending on which SOAT enzyme and its orientation, the cholesteryl ester product will be loaded into a very low-density lipoprotein particle or mobilized and stored in a lipid droplet [20]. This latter function has been recognized to play an adaptive role in alleviating membrane stress caused by excess free cholesterol, but incidentally it can also dramatically increase the concentration of cholesteryl ester in lipid droplets [2, 20]. Hence, in contrast to free cholesterol trafficking via membrane contact sites, cholesteryl esters stored in lipid droplets, which could be subsequently de-esterified to free cholesterol, may be the source of crystallized cholesterol. Here, we investigate the hypothesis that SOAT-mediated esterification of cholesterol is required for the formation of crystals in lipid droplets of cholesterol-loaded hepatocytes and liver. We also compare our findings to cholesterol loaded macrophage, as pathological cholesterol crystals have been more extensively investigated in this cell type.

## 2. MATERIALS AND METHODS

### 2.1. Cell culture and reagents

Human hepatoma 3B (Hep3B) cells (catalog #HB-8064) and RAW 264.7 cells (catalog #TIB-71) were purchased from American Type Culture Collection. Cells were cultured on 3.5 cm glass-bottom poly-D-lysine coated dishes from Mattek (catalog #P35GC-1.5-14-C) in Dulbecco’s Modified Eagle Medium (DMEM) from Gibco (catalog #11965-092) and supplemented with 10% cosmic calf serum from Cytiva Hyclone (catalog #SH30087.03) using a humidified incubator that was maintained at 37°C and a CO2 level of 5%. Sources of the reagents and consumables are as follows. SOAT inhibitors sandoz 58-035 (catalog #S9318), avasimibe (catalog #PZ0190) and K-604 (catalog #SML1837), methyl-ß- cyclodextrin (catalog #C4555), and cholesterol (catalog #C8667) were from Sigma-Aldrich. Dimethyl sulfoxide (catalog #BP231-1) and 96-well optical bottom, black walled microplates (catalog #1256670) were from Fisher. mTOR inhibitor rapamycin (catalog #1292) was from Tocris Bioscience. Low-density lipoprotein (LDL) and acetylated LDL (acLDL) was from Kalen Biomedical (catalog #770200-8 and #770201-4). Hoechst 33342 (catalog #62249), BODIPY 493/503 (catalog #D3922), LysoTracker Red DND-99 (catalog #L7528), Amplex Red Cholesterol Assay kit (catalog #A12216), propidium iodide (catalog #P21493), and Pierce bicinchoninic acid (BCA) protein assay kit (catalog #23225) were from ThermoFisher Scientific.

### 2.2. Preparation of methyl-ß-cyclodextrin (MßCD)-cholesterol complex

Dry cholesterol power (386.65 g/mol, 1.067 g/ml at 25°C) was solubilized in 100% ethanol at 70°C to a final concentration of 100 mM. 1 ml aliquots of the cholesterol solution were added to 42 mg/ml MßCD (1320 g/mol) in deionized water with constant heating at 70°C to a final concentration of 5 mM cholesterol. The cholesterol-MßCD complexed solution was cooled overnight and sterile filtered, and then stored at 4°C. We have shown this solution loads cells with cholesterol [22] and it will be referred to as the cholesterol treatment.

### 2.3. Imaging and detection of cholesterol crystals

4x10^5^ Hep3B cells or 1x10^5^ RAW 264.7 cells were seeded in 3.5 cm glass-bottom poly-D-lysine coated dishes and allowed to adhere overnight. To induce cholesterol crystallization, Hep3B cells were treated the following day with cholesterol or 500 µg/ml LDL and RAW 264.7 cells were treated with cholesterol or 50 µg/ml acLDL. Fresh media and treatments were provided every 24 hours. LDL treatment protocol for Figure 2F and 2G is as follows. 5x10^4^ Hep3B cells were seeded and treated for a total of 14 days as indicated and fresh treatments were provided every 3 days. Cells were trypsinized and re-seeded after 7 days to maintain optimal confluency. At the endpoint, media was removed and replaced with phosphate buffered saline (PBS) containing 5 µg/ml Hoechst 33342 and incubated at 37°C for 30 minutes to stain nuclei. Cholesterol crystal images of live cells were acquired under polarized light on an Olympus IX73 Inverted epifluorescence microscope with an Olympus DP73 camera at 20x magnification. Nuclei were imaged in parallel to determine cell number per field of view to normalize crystal area to cell number. Images from three distinct locations around dish were taken for representative sampling. Location of images were first determined using bright-field view to reduce potential observational bias that may occur by first viewing the degree of crystals. All images in the same channel within the same experiment were taken at identical exposures.

### 2.4. Quantification of cholesterol crystals in cultured cells

Cholesterol crystal quantification was performed using ImageJ software (https://imagej.nih.gov/ij/). First, raw crystal images were converted from RGB to 8-bit format, then the pixel threshold was set between 20-255 for cholesterol or LDL treated cells and between 15-255 for acLDL treated cells. Next, the image was measured and % crystal area recorded. To determine cell number per field of view, corresponding Hoechst 33342 nuclei images were converted from RGB to 16-bit format. The pixel threshold was set between 7-65535 for Hep3B nuclei and between 15 to 20-65535 for RAW 264.7 nuclei. A watershed was applied to the image to separate overlapping nuclei into individual objects. Next, particle size (pixel˄2) was set between 3000-infinity for Hep3B nuclei and 2000-infinity for RAW 264.7 nuclei with circularity set between 0.00-1.00. Particles were measured to determine cell number. Normalization of cholesterol crystal positive (crystal^+^) area to cell number was calculated by dividing the % crystal^+^ area per 100 cells in the field of view. The mean of three crystal images per plate was considered to represent one biological replicate and is shown in results as a single data point. Each biological replicate represents an independent experiment.

### 2.5. Quantification of cholesterol crystals in mouse liver sections

This was done as described previously [23]. Briefly, frozen liver sections were embedded in optimal cutting temperature compound (OCT), snap frozen on dry ice and cryo-sectioned at 10 mm thickness. All sections reached room temperature before being cover slipped with glycerol as mounting medium. Cholesterol crystals were immediately visualized under polarized light microscopy and analyzed with ImageJ software, as described in [10, 14].

### 2.6 Fluorescence imaging of lipid droplets

1.5x10^5^ Hep3B cells or 1x10^5^ RAW 264.7 cells were seeded in 3.5 cm glass-bottom poly-D-lysine coated dishes and allowed to adhere overnight. Cholesterol crystallization was induced by cholesterol treatment, as indicated in figure legends. At the endpoint, media was removed and replaced with serum-free DMEM containing 5 µg/ml Hoechst 33342 to stain nuclei (blue filter cube – ET DAPI), 2 µM BODIPY 493/503 to stain lipid droplets (green filter cube – FITC-A-Basic- OFF), and 50 nM LysoTracker Red DND-99 to stain lysosomes (red filter cube – TRITC-A-Basic-OFF). Cells were incubated at 37°C for 30 minutes. Prior to imaging, media was replaced with PBS. Fluorescence images of live cells were acquired alongside crystal images under polarized light on an Olympus IX73 Inverted epifluorescence microscope with an Olympus DP73 camera at 40x magnification. Images containing brightness/contrast enhancements were applied equally to all images within the figure and were not applied prior to image quantification.

### 2.7. Lipid extraction and cholesterol quantification

For Hep3B cells, 1x10^6^ were seeded in 3.5 cm dishes and allowed to adhere overnight. After treatment, cells were washed twice with cold PBS, collected in PBS by scraping, and centrifuged at 13,000 g for 10 minutes at 4°C. The pellets were then resuspended in hexane:isopropanol (3:2, v/v). For liver, approximately 50-100 mg frozen sample was homogenized in hexane:isopropanol (3:2, v/v). Afterward, samples were heated at 75°C for 5 minutes, and then centrifuged at 13,000 g for 10 minutes at room temperature. The soluble fraction containing lipid was transferred to a new tube and dried at 55°C overnight. Protein-containing pellets from Hep3B samples were dried, resuspended in 0.2 N sodium hydroxide and solubilized overnight at 37°C in order to normalize by protein content. Liver sample was normalized by tissue sample weight. The dried liver sample was resuspended in isopropanol:NP-40 (v/v, 9.9:0.1) and an aliquot of it was dried again at 55°C. This as well as the dried samples from cultured cells were resuspended in 300 µl of 1x Amplex Red Cholesterol Assay kit reaction buffer and diluted in 1x buffer, as necessary to get a reading within the standard curve. Cholesterol assay was performed according to manufacturer’s instructions. Briefly, 50 µl of working solution (containing 1x reaction buffer, 300 µM Amplex Red reagent, 2 U/ml horseradish peroxidase, 2 U/ml cholesterol oxidase, with or without 0.2 U/ml cholesterol esterase) was added to 50 µl lipid sample resuspended in 1x reaction buffer in a 96-well optical bottom, black walled microplate and then incubated at 37°C for 30 minutes, under protection from light. Fluorescence was measured using a Biotek Synergy HT microplate reader using excitation/emission at 530-560/590 nm. The concentration was determined by comparing to a standard curve. The free cholesterol quantity was determined by excluding cholesterol esterase in the assay, while total cholesterol was quantified by including cholesterol esterase. Esterified cholesterol was determined by subtracting free cholesterol from total cholesterol. Hep3B cellular protein content used for normalization was determined using the BCA protein assay.

### 2.8. Electron microscopy

1.2x10^6^ Hep3B cells treated as described in figure legend were resuspended in 700 μl of DMEM. FGP fixative solution (1.25% paraformaldehyde, 2.5% glutaraldehyde, 0.03% picric acid) was added to the suspension at a 1:1 ratio. Cells were fixed for 10 minutes at room temperature and then centrifuged at 2,000 rpm for 5 minutes at 4°C and then stored at 4°C. Samples were embedded in Epon resin and ultrathin sections were prepared using a Reichert Ultracut-S microtome and imaged with a Tecnai 12 (20-120 kV) transmission electron microscope equipped with a 2k x 2k CCD camera. The sample preparation and imaging were performed in the Arruda laboratory at UC Berkley.

### 2.9. Immunoblot

8x10^5^ Hep3B cells were seeded in 3.5 cm dishes and allowed to adhere overnight. After appropriate treatment, cells were washed twice with cold PBS and lysed in RIPA buffer (150 mM NaCl, 50mM Tris-HCl pH 7.5, 0.1% w/v sodium dodecyl sulfate (SDS), 0.5% w/v Na-Deoxycholate, 1% v/v Nonidet p40, 1 mM EDTA, 1 mM EGTA, 2.5 mM Na pyrophosphate, 1 mM NaVO4, 10 mM NaF) supplemented with 1x protease inhibitor cocktail (catalog #11873580001) and 1x phosphatase inhibitor cocktail (catalog #P5726) from Sigma-Aldrich, and 1% 2-mercaptoethanol. Protein concentration was determined using the Pierce BCA protein assay kit. 10 μg protein was loaded onto Bolt^TM^ 4-12% Bis-Tris mini protein gels (catalog #NW04125BOX) or Novex^TM^ 14% Tris-Glycine mini protein gels (catalog #XP00145BOX) from Invitrogen. Electrophoresis was run at neutral pH in NuPAGE^TM^ MOPS running buffer (catalog #NP0001) or Novex Tris-Glycine SDS running buffer (catalog #LC2675). Proteins were transferred to nitrocellulose membrane from Bio-Rad (catalog #1620115), blocked in tris-buffered saline containing 0.1% tween 20 (TBST) and 5% powdered skim milk, and incubated overnight at 4°C with primary antibody detecting light chain 3 type 1 and 2 (Cell Signaling technology, catalog #3868), SOAT1 (catalog #84476), and glyceraldehyde 3 phosphate dehydrogenase (Cell Signaling technology, catalog #2118). After wash, anti-rabbit HRP-conjugated antibody (Cell signaling technology, catalog #7074S) was incubated 1/5000 in TBST containing 5% milk at room temperature for 1 hour. Membranes were washed again and then incubated with Super Signal West Femto Maximum Sensitivity Substrate from ThermoFisher Scientific (catalog #34096) to generate chemiluminescence that was detected by a BioRad ChemiDoc MP Imaging System and analyzed using ImageJ software.

### 2.10. RNA isolation and gene expression

After appropriate treatment, media was removed, and cells were lysed and collected in 500 μl TRIzol Reagent from ThermoFisher Scientific (catalog #15596018) and total RNA was isolated according to manufacturer’s protocol. 500 ng of purified RNA was used for cDNA synthesis using Maxima H Minus First Strand cDNA Synthesis Kit with dsDNase from ThermoFisher Scientific (catalog #K1672) according to protocol. Quantitative polymerase chain reaction (qPCR) was performed on a Bio-Rad CFX384 Real-Time PCR Detection System using PowerUP^TM^ SYBR^TM^ Green Master Mix from ThermoFisher Scientific (catalog #A25742). Cycle thresholds were normalized to glyceraldehyde-3-phosphate dehydrogenase levels and displayed as relative mRNA level.

### 2.11. CRISPR/cas9-based gene editing of Hep3B cells

The stable cas9 expressing Hep3B (Hep3B^cas9^) cell line was generated via transfection with Cas9 plasmid (Sigma catalog #CAS9HYGROVP) at a ratio of 6:1 using lipofectamine (Roche, catalog #6365787001), following manufacturer’s instructions. Cells were selected in medium containing 8 μg/ml blasticidin (Sigma, catalog# 15205-25MG). Single clones were isolated using a cloning cylinder (Sigma, catalog #TR-1004) and maintained on 2 μg/ml blasticidin. To validate active Cas9 expression, clones were transfected with a synthetic guide RNA (Invitrogen, catalog# A355533) and cultured for two days, after which then genomic DNA was isolated for detection of gene editing efficiency using the GeneArt genomic cleavage assay kit (Invitrogen, catalog #A24372). Hep3B^cas9^ cells had high cleavage efficiency and thus was chosen for generating knockout cell lines using CRISPR. To generate SOAT1 or SOAT2 knockout cell lines, 9 x 10^5^ of Hep3B^cas9^ were seeded onto 60 mm culture dish, CRISPR guide RNA lentivirus targeting human SOAT1 (Invitrogen, Assay ID: CRISPR693296LV and CRISPR693330LV) or SOAT2 (Invitrogen, Assay ID: CRISPR824991LV and CRISPR824999LV) were added to the cell culture at a multiplicity of infection (MOI) of 5. All lentiviral knockouts were selected by culturing cells in the presence of puromycin (3 μg/ml) 48 hours after infection.

### 2.12. Animal study and liver collection

This was done as previous [23]. Animal handling and procedures performed were approved by the University of Saskatchewan’s Animal Care Committee. Male and female mice were housed at 21°C on a 12-hour light/dark cycle and provided ad libitum access to food and water. C57bl/6J mice were purchased from The Jackson Laboratory (www.jax.org/strain/000664). Mice with flox alleles for the NRF1 and NRF2 gene (*nfe2l1^flox/flox^*; *nfe2l2^flox/flox^*) were generated as previously described [22, 23]. To delete NRF1 and NRF2 in hepatocytes, recombination of flox alleles to remove respective gene elements was done by retroorbital infection of mice, while under isoflurane anesthesia, with 2.0 x 10^11^ particles of liver targeting serotype 8 adeno-associated virus expressing Cre recombinase via hepatocyte-specific thyroxine binding globulin promoter (AAV-CRE), as previous [23]. Littermate controls received virus expressing green fluorescent protein (AAV-GFP). AAV-CRE (AAV8.TBG.PI.Cre.rBG) and AAV-GFP (AAV8.TBG.PI.eGFP.WPRE.bGH) viruses were acquired from the University of Pennsylvania Vector Biocore. Mice continued feeding on regular chow (Prolab RMH 3000 from LabDiet) for 7 days post-infection. Then, mice continued on regular control chow or were fed MASH inducing high fat, fructose, and cholesterol (HFFC) diet (Research diets, catalog# D09100310i), which contains 40% fat (75% palm oil/11% lard/14% soybean oil), 10% sucrose, 20% fructose and 2% cholesterol, as indicated in figure 6A. At the same time, mice were treated with daily intraperitoneal injections of Avasimibe (5mg/kg) or vehicle (10 % DMSO in PBS) for 10 days. In longer term 24-week HFFC diet study, wild type C57bl/6J mice were fed and treated as illustrated in figure 6G. At the endpoint of studies, mice were euthanized under anesthesia with 3% isoflurane at an oxygen flow rate of 1 L/minute. Vasculature of anesthetized mice was flushed with 20 ml phosphate buffered saline that was at room temperature. A portion of liver was snap frozen and stored at -80 °C. A second portion from left lobe was fixed in 10 % neutral buffered formalin, embedded in wax, sectioned at 5 µm thickness, and stained with Hematoxylin and Eosin (H&E). A third portion was embedded in OCT compound, snap frozen on dry ice, and cryo-sectioned at a 10 μm thickness, and then was used for crystal detection (described in section 2.5).

### 2.13. Statistical analysis

All data analysis was performed using GraphPad PRISM (version 9.5.1). Significance was defined as p<0.05 using t-test, simple regression analysis, and one-way or two-way analysis of variance (ANOVA) with Dunnett’s or Tukey’s post-hoc test, as indicated in figure legends. Data are expressed as mean ± standard error of the mean (SEM). Sample size is indicated in each figure. A minimum of three biological replicates (independent experiments) were used for each study.

## 3. RESULTS

### 3.1. Time- and dose-dependent crystal formation in lipid droplets of cholesterol loaded Hep3B cells

We reasoned that a cell culture model would be the most efficacious approach for investigating the mechanism underlying hepatocyte cholesterol crystal formation. However, while such crystals have been extensively studied in macrophage and its relationship to atherosclerosis [17, 18, 24, 25], to our knowledge only two studies have been undertaken on cultured hepatocytes [10, 14]. In these works, cultured HepG2 cells were treated with LDL plus oleic acid, which led to a modest level of lipid droplet crystals by day 20. As this paradigm had limited effect and required a relatively long treatment period, we attempted to establish a more rapid and robust cell culture model. For this, we opted to use human hepatocyte Hep3B cells, rather than HepG2 cells, because they formed more uniform monolayers that are better suited for imaging. Hep3B cells were loaded with cholesterol upon incubation with media containing methyl-ß- cyclodextrin (MßCD) that was in a saturated complex with cholesterol (referred to as cholesterol treatment). As described [22], this treatment results in the transfer of cholesterol in this complex to the cell membrane and thus loading the plasma membrane with free cholesterol. Cells can adapt by redistributing cholesterol, increasing its export, and decreasing lipoprotein uptake and cholesterol synthesis as well as by esterifying cholesterol and storing it in lipid droplets, a process mediated by SOAT activity [19, 21].

The formation of cholesterol crystals in cells was detected via its birefringent property that can be imaged using polarizing light microscopy, as described in [8, 10, 14]. The measured crystal area per field was normalized by cell number, which was determined via fluorescence detection of Hoechst 33342 stained nuclei. Using this approach, we first performed a time course study by treating Hep3B cells with 200 µM cholesterol for 6-96 hours. Cholesterol treated cells had a robust level of crystals by the 48- and 96-hour timepoint, whereas controls had no detectable crystals (Figures 1A and 1B). We also assessed the dose-effect by treating cells with 0-200 µM cholesterol for 48 hours. Cells treated with 100 µM and 200 µM cholesterol, but not 50 µM, developed a significant level of crystals (Figures 1C and 1D). Interestingly, biochemical analysis revealed cellular cholesteryl ester levels positively correlated with crystal abundance, with a higher correlation coefficient than free cholesterol level (Figures S1A and S1B). Moreover, in a resolution experiment, we found crystals induced by cholesterol treatment resolved by approximately 50% after switching the cells back to control media for 48 hours (Figures S1C and S1D). Next, we examined whether the crystals localized to lipid droplets or other sub-cellular regions such as lysosomes. Hep3B cells were treated with 200 µM cholesterol or control media for 48 hours. Images were acquired like in Figures 1A and 1C, except at a higher magnification and while also imaging BODIPY-stained lipid droplets and Lysotracker Red-labelled lysosomes (Figure 1E). Crystals were almost exclusively detected at the periphery to nearly all lipid droplets, in a manner like that described for MASH-associated crystals [10, 14]. Altogether, these results show cholesterol loading of Hep3B cells, via a MßCD-cholesterol complex, results in a robust time- and dose-dependent induction of lipid droplet localized crystals that is reversible and correlates with cholesteryl ester content. Hence, we deemed it well suited for mechanistic investigation.

**Figure 1.**
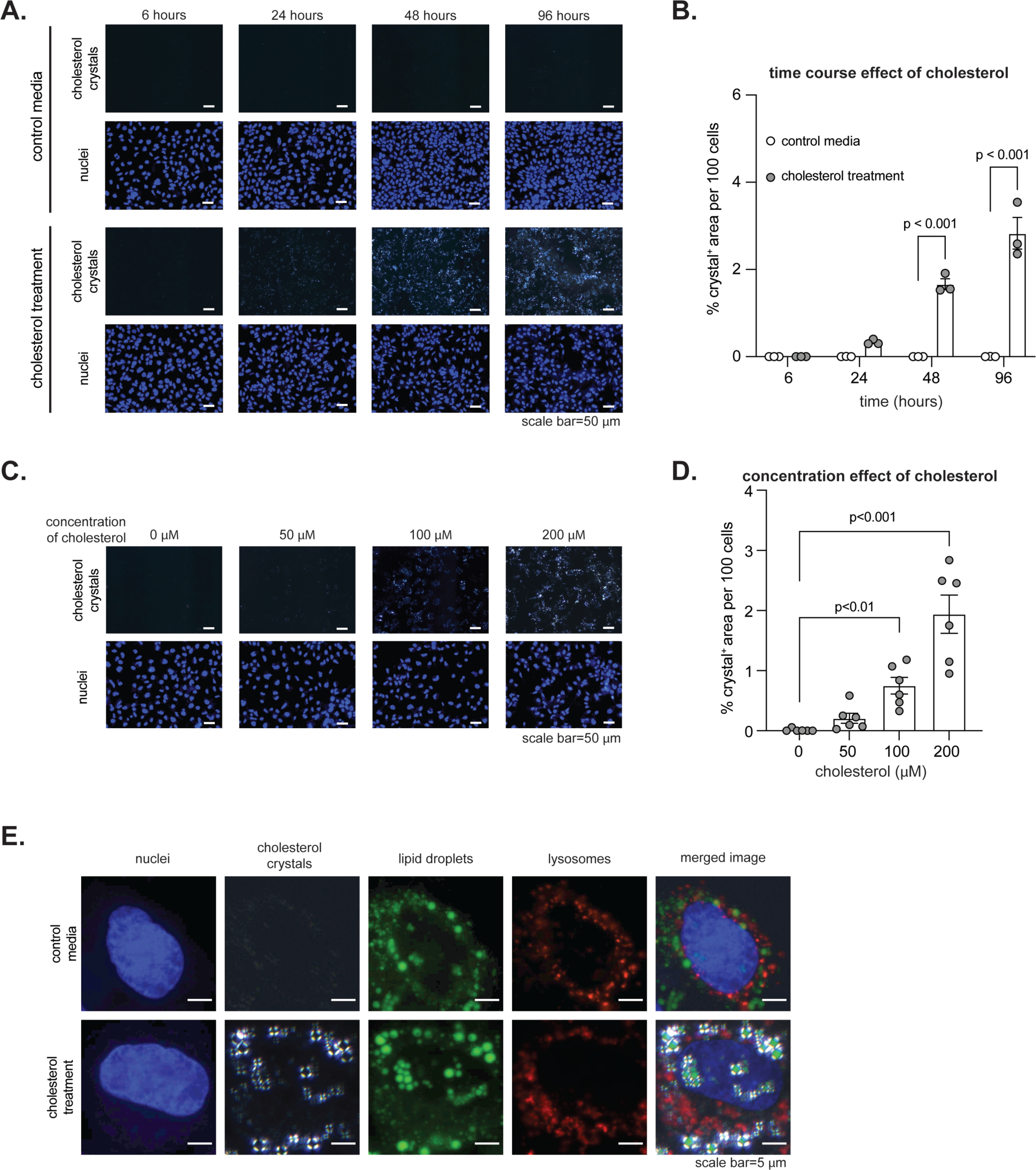
Time- and dose-dependent crystal formation in lipid droplets of cholesterol loaded Hep3B cells. (A) Representative images of Hep3B cells treated with or without 200 µM cholesterol for 6-96 hours. Detected crystals and nuclei is indicated in panel. (B) Quantified cholesterol crystals in (A) to examine time effect on crystal formation (n=3). The p-value was determined by t-test, adjusted for multiple comparison. (C) Representative images of Hep3B cells treated with 0-200 µM cholesterol for 48 hours. (D) Quantified cholesterol crystals in (C) to examine cholesterol concentration effect on crystal formation (n=6). The p- value was determined by one-way analysis of variance, with Dunnett’s post-hoc test. (E) Representative images of Hep3B cells treated and stained as indicated for 48 hours to examine the subcellular localization of cholesterol crystals. Scale bar for (A), (C), and (E) is shown in panel. Data in (B) and (D) are mean ± standard error of the mean, with individual data points shown.

**Figure 2.**
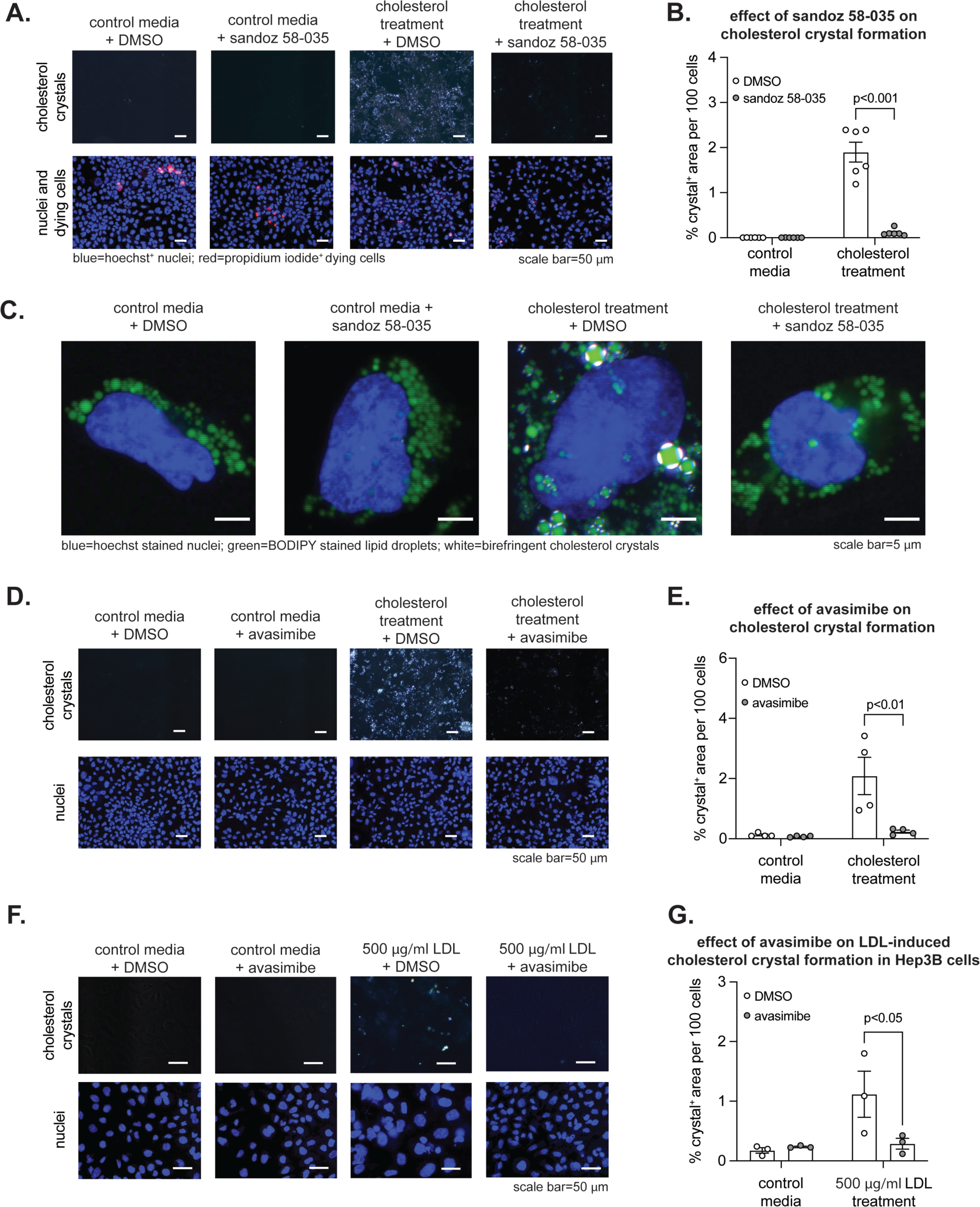
SOAT inhibitor prevents crystal formation in cholesterol loaded Hep3B cells. (A) Representative images of cells treated with or without cholesterol ± 1 µg/ml sandoz 58-035 for 48 hours. (B) Quantified cholesterol crystals in (A) to examine the effect of sandoz 58-035 on crystal formation (n=6). (C) Representative images of cells treated and stained as indicated. (D) Representative images of cells treated with or without cholesterol ± 10 µM avasimibe. (E) Quantified cholesterol crystals in (D) to examine the effect of avasmibe on crystal formation (n=4). (F) Representative images of cells treated with or without 500 µg/ml low density lipoprotein (LDL) ± 10 µM avasimibe. (G) Quantified cholesterol crystals in (F) to examine the effect of avasmibe on crystal formation (n=3). The p-value in (B), (E), and (G) was determined by t-test, adjusted for multiple comparison. Scale bar for (A), (C), (D), and (F) is shown in panel. Data in (B), (E), and (F) are mean ± standard error of the mean, with individual data points shown.

### 3.2 SOAT inhibitor prevents crystal formation in Hep3B cells

To investigate whether esterification by SOAT activity is required for cholesterol crystals to form in lipid droplets, we examined the effect of well-established SOAT inhibiting compound sandoz 58-035 [26, 27]. Hep3B cells were treated with cholesterol or control media in the presence of 1 µg/ml 7andoz 58- 035 or vehicle (dimethyl sulfoxide; DMSO). Sandoz 58-035 mediated SOAT inhibition was confirmed by a lack of cholesteryl ester accumulation in cholesterol treated cells (Figure S2A). Most remarkable however was that sandoz 58-035 almost completely blunted formation of crystals (Figures 2A and 2B). Propidium iodide staining of dying cells ruled out that this effect was due to altered cell viability, as it was not different between cells receiving sandoz 58-035 or vehicle (Figures 2A and S2B). Conversely, sandoz 58-035 did cause the anticipated changes in expression of liver X receptor and sterol response element binding protein 2 target genes (Figure S2C), a result consistent with increased free cholesterol. Higher magnification images showed that sandoz 58-035 blocked formation of lipid droplet localized crystals (Figure 2C). To confirm this effect was due to inhibiting SOAT activity, we performed a study similar to Figure 2A except in place of sandoz 58-035 we used avasimibe, a chemically distinct but also well-established SOAT inhibitor that has been used in clinical investigations [28]. Like sandoz 58-035, avasimibe dramatically reduced crystal formation in cholesterol treated Hep3B cells (Figures 2D and 2E). Moreover, to ensure our findings were not specific to the mode by which Hep3B cells were loaded with cholesterol we also performed a study similar to Figure 2D except in place of MßCD-cholesterol complex we used 500 µg/ml LDL treatment for 14 days to load cells with cholesterol, a model similar to that used by Ioannou and colleagues with HepG2 cells [10, 14]. Though the effect was less robust than MßCD-cholesterol complex, LDL induced significant crystal formation and, this effect was blunted by avasimibe (Figures 2F and 2G). Altogether, we show SOAT-mediated cholesteryl esterification underlies formation of lipid droplet crystals in hepatocytes.

### 3.3. SOAT inhibition blocks formation of crystal cleft deposits, independent of autophagy

A common feature of cholesterol crystals in atherosclerotic plaques is crystalline lattice deposits, identified as plate-like clefts under election microscope imaging, due to solvent effects during processing [24, 25, 29, 30]. It is not known whether cholesterol overload induces similar clefts in hepatocytes. We investigated whether lipid droplet crystals in Hep3B cells may serve as a nucleating site for crystal lattice deposits. To do this, we used transmission election microscope imaging on Hep3B cells treated with cholesterol or control media combined with or without sandoz 58-035. As expected, cells treated with control media did not have any clefts. However, structures resembling crystal clefts were identified in a fraction (15-20%) of cholesterol + DMSO treated cells, but not in any of the cells treated with cholesterol + sandoz 58-035 (Figure 3A). Another notable feature in cholesterol treated cells was increased autophagy, irrespective of sandoz 58-035, as recognized by the high frequency of autophagosomes (Figure 3A) and immunoblot detection of autophagy marker, light chain 3 type II levels (Figures S3A and S3B). As autophagy could promote crystal dissolution, we tested whether enhancing autophagy activity using rapamycin reduced the level of crystals in cholesterol treated cells. However, while rapamycin induced autophagy, it was modest compared to cholesterol treatment (Figures 3B and 3C) and had no effect on crystal levels (Figures 3D and 3E). These findings indicate lipid droplet localized crystals may serve as a nucleating site for the development of crystal lattices, independent of autophagy, and that this effect is blocked by SOAT inhibition.

**Figure 3.**
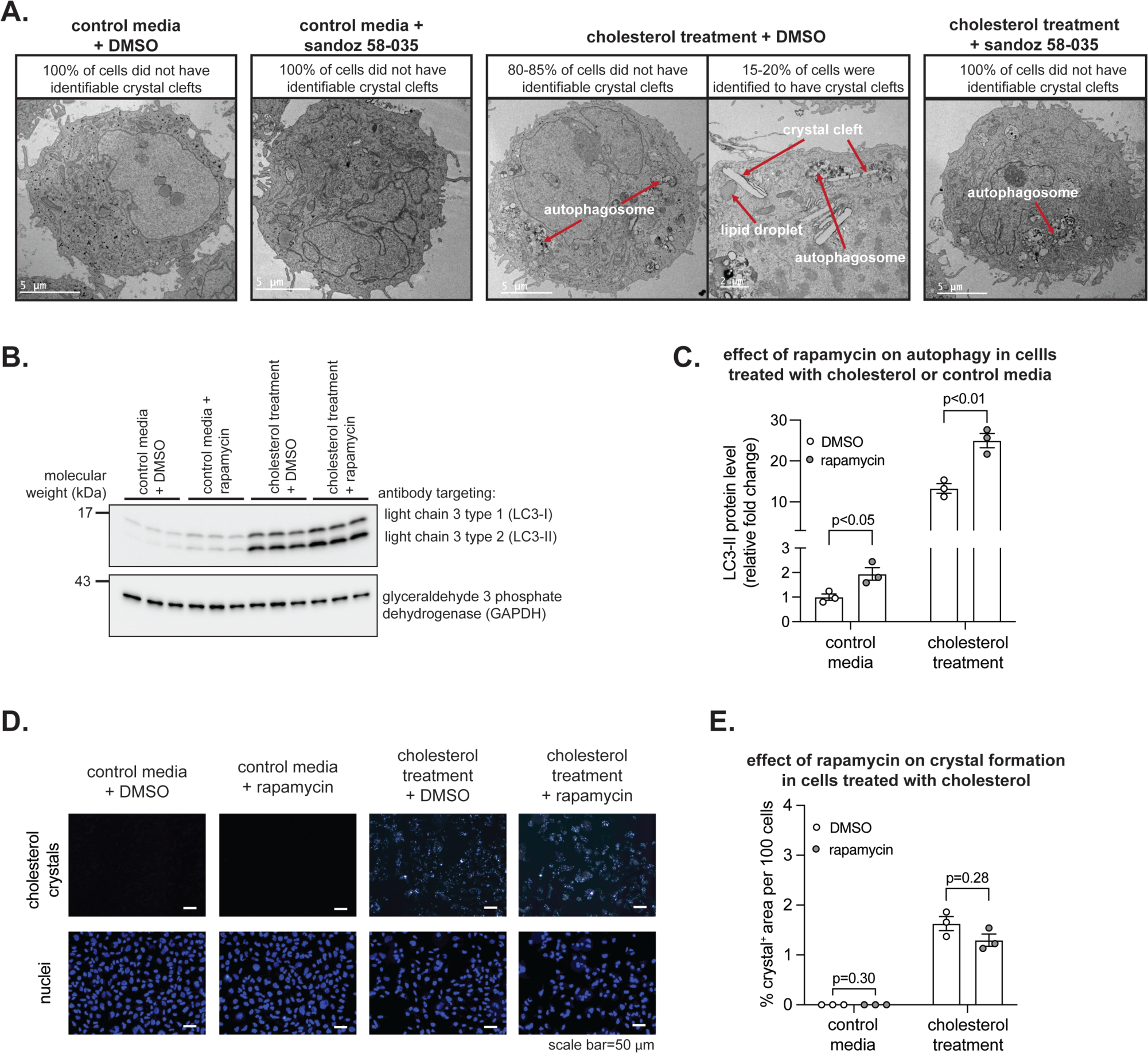
SOAT inhibition blocks formation of crystal cleft deposits, independent of autophagy. (A) Representative transmission electron microscope images of Hep3B cells treated with or without cholesterol ± 1 µg/ml sandoz 58-035. The number of cells examined for each group is indicated in the panel. (B) Immunoblot of indicated proteins extracted from Hep3B cells treated with or without cholesterol ± rapamycin. (C) Quantified immunoblot in (C) to examine the effect of rapamycin on autophagy (n=3). (D) Representative images of cells treated with or without cholesterol ± 100 nM rapamycin. (E) Quantified cholesterol crystals in (D) to examine the effect of rapamycin-induced autophagy on crystal formation (n=3). Scale bar for (A) and (D) is shown in panel. Red arrows in (A) highlight the indicated cellular features. Data in (C) and (E) are mean ± standard error of the mean, with individual data points shown.

### 3.4. SOAT1 mediates crystal formation in Hep3B cells

While our results using sandoz 58-035 and avasimibe show SOAT activity is required for crystal formation, it does not discern whether this was due to SOAT1 or SOAT2, as these agents can inhibit both [27, 28]. But since SOAT1 is primarily recognized to produce cholesteryl ester that is stored in lipid droplets [20, 21], we reasoned that SOAT1 impairment may be sufficient to blunt cholesterol crystal formation. We initially tested this by treating cells with a 0-30 nM concentration range of K-604, a SOAT1-selective inhibitor that has greater than 200-fold selectivity for SOAT1 and a half maximal inhibitory concentration of 68 nM [31]. In cholesterol treated Hep3B cells 30, 10, and 3 nM K-604 blunted crystal formation, an effect that did not occur at lower K-604 concentrations (Figures 4A and 4B). This result is consistent with a SOAT1 specific role in lipid droplet crystals. To confirm, we examined SOAT1 deficient (*SOAT1^-/-^*) cells (Figure 4C), generated using clustered regularly interspaced short palindromic repeat (CRISPR)-based genome editing on CRISPR associated 9 (cas9) expressing Hep3B (Hep3B^cas9^) cells. Cholesterol treatment caused crystal formation in Hep3B^cas9^ control cells, and this effect was strongly blunted in two independent *SOAT1^-/-^* cell lines (Figures 4D and 4E). Conversely, examination of *SOAT2^-/-^* cells in a similarly designed experiment revealed SOAT2 is dispensable for crystal formation (Figures S4A and S4B). Thus, our findings show that SOAT1 mediates the formation of lipid droplet cholesterol crystals.

**Figure 4.**
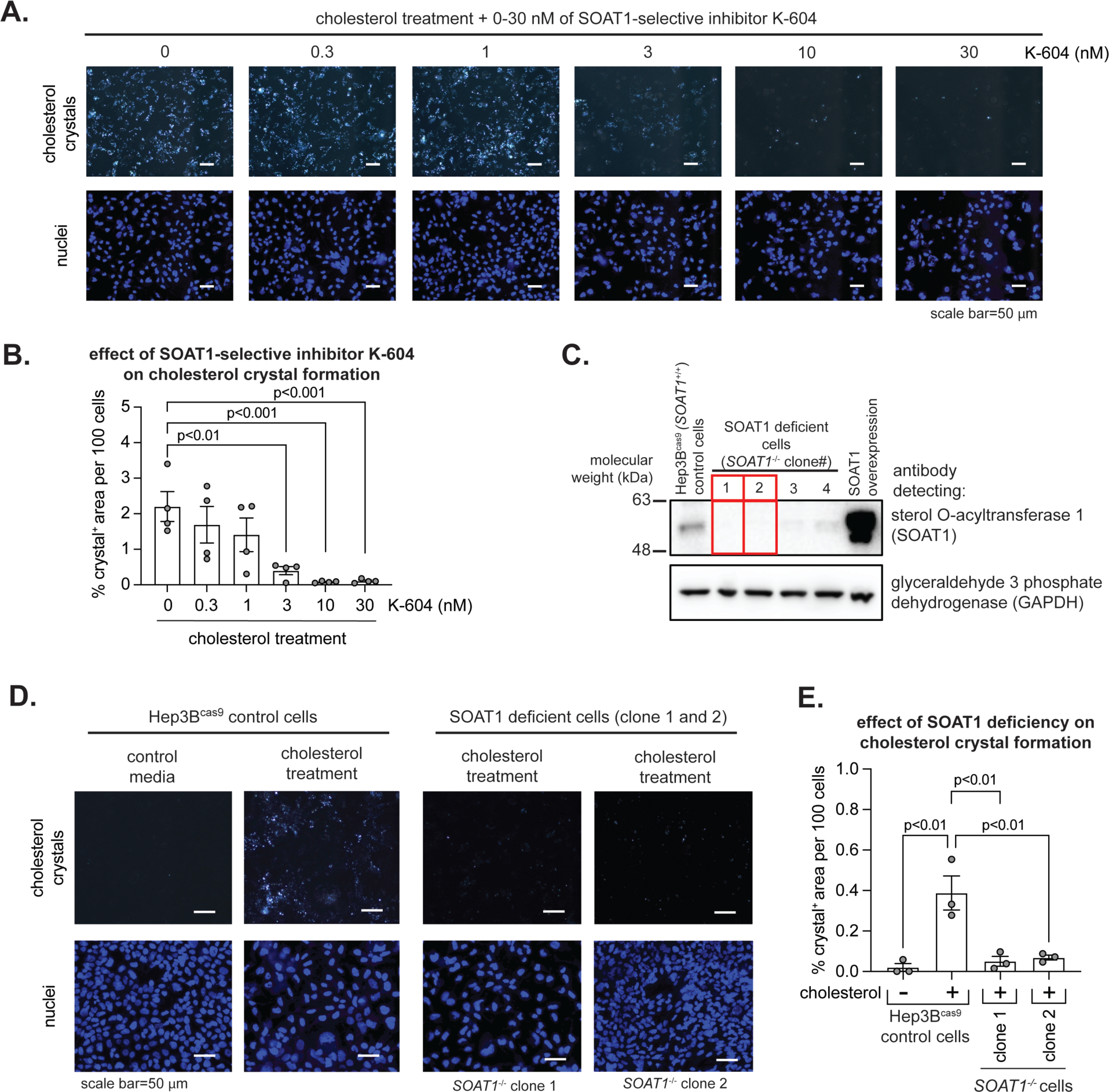
SOAT1 mediates crystal formation in cholesterol loaded Hep3B cells. (A) Representative images of cells treated with cholesterol + 0-30 nM SOAT1 selective inhibitor, K-604. (B) Quantified cholesterol crystals in (A) to examine the effect of K-604 on crystal formation (n=4). (C) Immunoblot for SOAT1 and GAPDH loading control of indicated proteins extracted from control Hep3B^cas9^ and *SOAT1* CRISPR-edited cells as well as Hek 293 cells over-expressing human SOAT1 (far right lane), which served as a positive control. (D) Representative images of Hep3B^cas9^ cells and SOAT1 deficient cells treated as indicated with or without cholesterol. (E) Quantified cholesterol crystals in (D) to examine the effect of SOAT1 deficiency on crystal formation (n=3). The p-value in (B) and (E) was determined by one-way analysis of variance, with Dunnett’s post-hoc test. Scale bar for (A) and (E) is shown in panel. Data in (B) and (E) are mean ± standard error of the mean, with individual data points shown.

### 3.5. SOAT inhibition prevents crystal formation in RAW 264.7 macrophage

Cholesterol crystals have been investigated extensively in macrophage [17, 18, 24, 25], but to our knowledge these works have not indicated that SOAT activity contributes to the formation of cholesterol crystals. To assess whether there may be such a role for SOAT activity in the macrophage, we examined the effect of SOAT inhibition using sandoz 58-035 on the RAW 264.7 macrophage cell line. RAW 264.7 cells were loaded with cholesterol by treating with 200 µM cholesterol, as described for Figure 1, or with 50 µg/ml acetylated LDL (acLDL). Cholesterol treatment (Figures S5A and S5B) and acLDL (Figures S5C and S5D) caused substantial formation of cholesterol crystals by 24- and 48-hour timepoints. Moreover, crystals in cholesterol treated RAW 264.7 cells were associated with lipid droplets and not with the lysosome (Figure 5A). And just as occurred in Hep3B cells, SOAT inhibition with sandoz 58-035 blocked the formation of crystals in cells treated with acLDL (Figures 5B and 5C) and with cholesterol (Figure S5E). Hence, our findings show SOAT activity mediates formation of lipid droplet crystals in macrophage.

**Figure 5.**
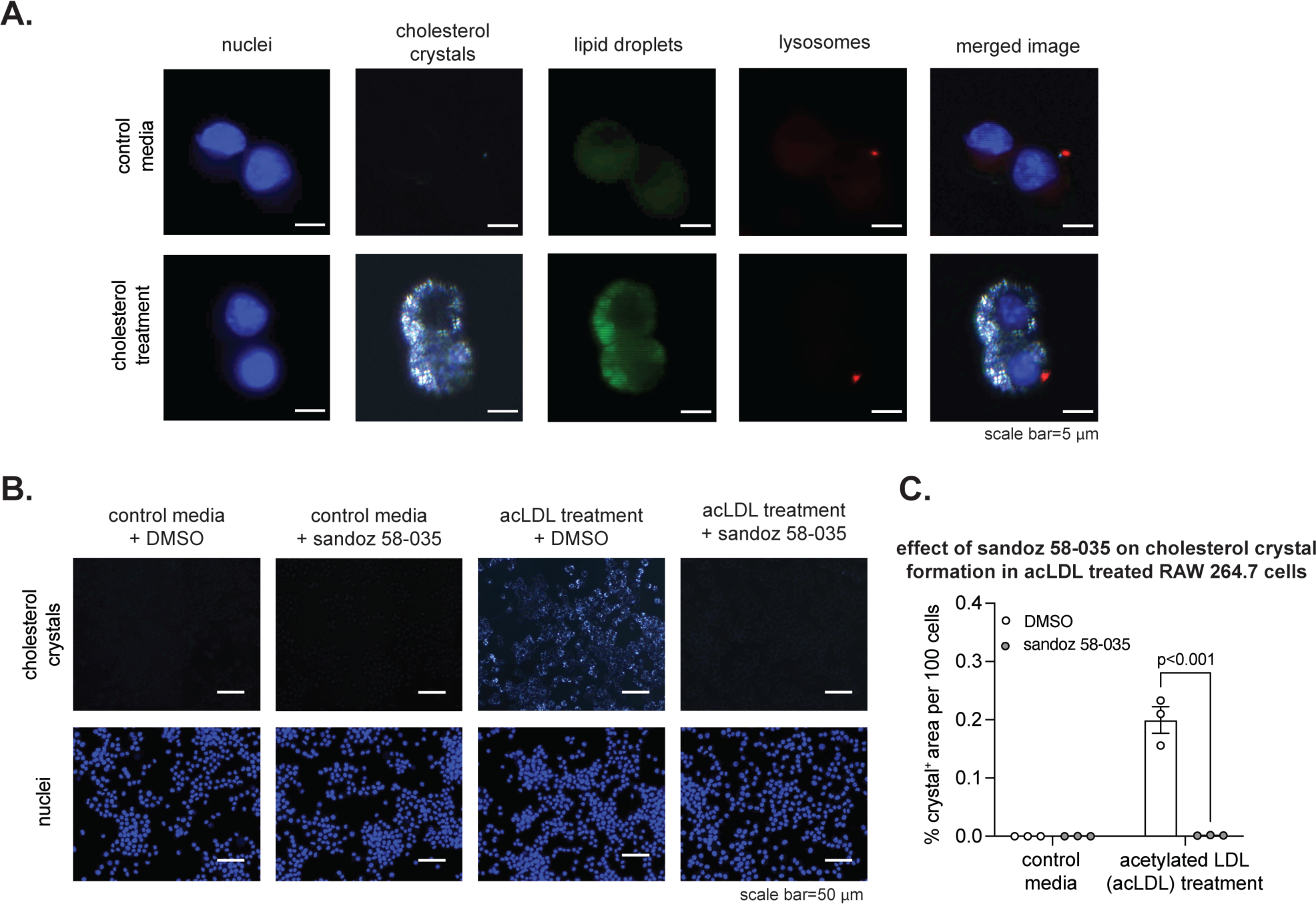
SOAT inhibition prevents crystal formation in cholesterol loaded RAW 264.7 macrophage. (A) Representative images of RAW 264.7 cells treated with or without cholesterol to examine the effect on crystal formation and its subcellular localization to lipid droplets. (B) Representative images of cells treated with or without 50 µg/ml acetylated LDL (acLDL) ± 1 µg/ml sandoz 58-035 for 48 hours. (C) Quantified cholesterol crystals in (B) to examine effect of sandoz 58-035 on crystal formation (n=3). The p-value was determined by t-test, adjusted for multiple comparison. Scale bar for (A) and (C) is shown in panel. Data in (C) are mean ± standard error of the mean, with individual data points shown.

**Figure 6.**
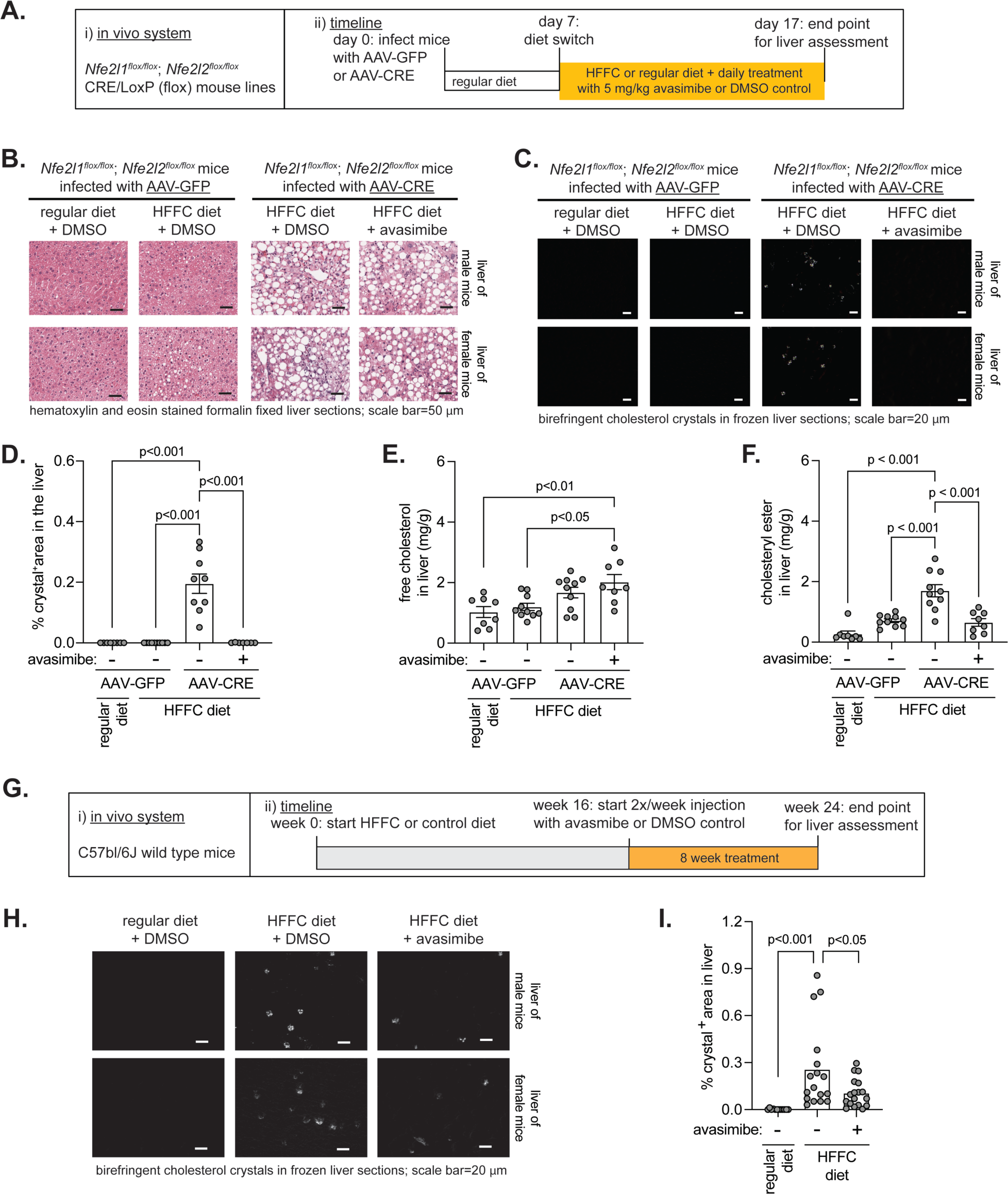
SOAT inhibition blocks crystal formation in mice with hepatic cholesterol overload. (A) Schematic of the experiment designed to test whether avasimibe prevents crystal formation in liver of mice with hepatic cholesterol overload. (B and C) Representative images of hematoxylin and eosin-stained fixed liver sections (B) or crystal detection in frozen liver sections (C) from male and female mice with or without hepatocyte Nrf1 and Nrf2 deficiency and fed cholesterol enriched or regular diet. (D-F) Quantified cholesterol crystals (D), free cholesterol (E), and cholesteryl ester (F) in liver to examine the effect of avasimibe (n=3-5 males and 4-6 females). (G) Schematic of the experiment designed to test whether avasimibe treatment between week 16-24 reduces crystal formation in liver of wild mice fed HFFC diet for 24 weeks. (H) Representative images of crystal detection in frozen liver sections from male and female mice fed HFFC or regular diet with or without twice per week avasimibe injections for 8 weeks. I) Quantified cholesterol crystals in liver corresponding to (H) to examine the effect of avasimibe in mice fed HFFC diet for 24 weeks. (n=6-8 males and 6-10 females). The p-value in (D-F and I) was determined by one-way analysis of variance, with Tukey’s post-hoc test. Scale bar for B, C, and H are shown in panel. Data in (D-F and I) are mean ± standard error of the mean, with individual data points shown.

### 3.6. SOAT inhibition blocks crystal formation in mice with hepatic cholesterol overload

We then used mice to investigate whether *in vivo* SOAT activity contributes to hepatocyte cholesterol crystals. In previous work using C57bl/6J mice fed a high fat plus cholesterol enriched diet, hepatic crystals were identified after 24-30 weeks of diet exposure [8, 10]. We recently showed that mice with combined hepatocyte deficiency of transcription factors Nrf1 (*Nfe2l1*) and Nrf2 (*Nfe2l2*) develop hepatic crystals after only 3 weeks of the diet [23]. Here, we first took advantage of this rapid crystal onset model. Liver of male and female mice with flox alleles in the genes encoding Nrf1 and Nrf2 (*Nfe2l1^flox/flox^*; *Nfe2l2^flox/flox^*) were infected with adeno-associated virus expressing cre recombinase (AAV-CRE) or green fluorescent protein (AAV-GFP) under control of a hepatocyte specific promoter (Figure 6A). Seven days later mice were fed a diet high in fat, fructose, and cholesterol (HFFC), as previous [23], or continued on a regular diet while also receiving daily intraperitoneal injections of DMSO vehicle or 5 mg/kg avasimibe. After 10 days of feeding, liver was collected for analysis. Hematoxylin and eosin staining of liver sections revealed dramatic lipid droplet accumulation in AAV-CRE infected mice fed HFFC diet (Figure 6B), consistent with previous [23]. As expected, AAV-GFP infected mice fed HFFC or regular diet had no detectable crystals (Figures 6C and 6D). Conversely, AAV-CRE infected mice fed HFFC diet and administered DMSO developed significant hepatic crystals (Figures 6C and 6D) as well as a trending increase in hepatic free cholesterol and increased hepatic cholesteryl ester (Figures 6E and 6F). In contrast, AAV-CRE infected mice fed HFFC diet and administered avasimibe had virtually no detectable crystals (Figures 6C and 6D), while having increased hepatic free cholesterol (Figure 6E) and decreased hepatic cholesteryl ester (Figure 6F). Finally, we investigated whether *in vivo* SOAT activity contributes to hepatocyte cholesterol crystals in wild type mice with prolonged 24-week exposure to MASH promoting HFFC diet (Figure 6G). Mice were fed HFFC diet or regular diet for 16 weeks and then received intraperitoneal injections of DMSO vehicle or 5 mg/kg avasimibe twice per week for the remaining 8 weeks of diet feeding. As expected, mice fed regular diet did not hepatic crystals, whereas mice fed the HFFC diet and administered DMSO had a significant level of hepatic crystals (Figures 6H and 6I). In contrast, hepatic crystals were significantly less in mice fed HFFC diet and administered avasimibe and were not significantly different than crystal level in regular diet fed mice (Figures 6H and 6I). Altogether, these results show blocking cholesterol esterification by inhibiting SOAT activity reduces cholesterol crystal formation in cholesterol overloaded liver.

## 4. DISCUSSION

Metabolic dysfunction-associated steatohepatitis (MASH) has become a global health concern [32, 33], but there is incomplete understanding of its pathogenesis and treatment. Hepatocyte cholesterol crystals are associated with MASH, but not simple steatosis, in humans and in rodent disease models that are induced by cholesterol enriched diet [3, 4]. These crystals may induce liver inflammation similarly to how they induce inflammation in atherosclerotic plaques [3, 17, 24, 29]. However, more study is needed to clarify if this is the case, since recent evidence shows the crystal sensing NLR family pyrin domain containing 3 (NLRP3) inflammasome is dispensable for cholesterol-associated MASH in mice [34] whereas NLRP3 has been shown to mediate inflammation in atherosclerotic plaques [17]. Investigating the mechanism by which hepatic cholesterol crystals form may reveal whether such crystals trigger liver inflammation and, if they do, which molecules could be targeted to alleviate MASH.

In this study, we sought to identify molecules that mediate hepatic cholesterol crystal formation. This was largely done using cholesterol loaded Hep3B cells, which we demonstrated to be a suitable, convenient experimental model. We discovered that inhibiting sterol O-acyltransferase (SOAT) activity in cholesterol loaded cells with sandoz 58-035 or avasimibe almost completely prevented crystal formation, and that this effect was not due to altering autophagy or cell viability. Using transmission electron microscopy (TEM), we identified cholesterol loaded cells with crystal lattice deposits, which, importantly, did not occur in cells treated with SOAT inhibitor, sandoz 58-035. We speculate that they may manifest stochastically when lipid droplet crystals nucleate. Moreover, using a selective inhibitor, K-604, or gene deficient cells, we showed that SOAT1 is the primary cholesteryl ester producing enzyme mediating crystal formation. And taking advantage of a transgenic mouse model that rapidly develops hepatic cholesterol crystals [23] and wild type mice fed HFFC diet for 24 weeks, we also showed inhibiting *in vivo* SOAT activity can prevent formation of hepatic cholesterol crystals. Altogether, our evidence supports our hypothesis that hepatic cholesterol esterification by SOAT1 is required to form cholesterol crystals in hepatocyte lipid droplets.

Based on our results combined with its well-established functions [20, 21], we suspect that SOAT1 promotes crystal formation by concentrating cholesteryl ester in lipid droplets. This raises the question as to whether the detected crystals were cholesteryl ester or de-esterified cholesterol. While our study does not discern which, de-esterified lipid droplet cholesterol seems a more likely scenario since cholesterol, not cholesteryl ester, has been shown to nucleate and form crystal deposits in cells [17, 24, 25, 29], like those we identified using TEM. Which enzyme primarily mediates cholesteryl ester hydrolase activity in hepatocytes is not known. Further investigation is needed to identify it and determine whether blocking its activity prevents hepatic crystal formation. Likewise, it remains unclear to what degree, if any, hepatic cholesterol crystals promote MASH and, if so, which molecular pathway senses and mediates the inflammatory signal. We anticipate such insight will reveal valuable targets for treating MASH.

There is emerging interest in targeting SOAT1, also known as acyl co-A cholesterol acyltransferase 1 (ACAT1), for neurological disorders [35, 36] and recent work has shown that avasimibe may alleviate hepatocellular carcinoma [37, 38]. Hence, considering our finding in this study, SOAT1 may be a plausible target for MASH and its progression. Interestingly, we also found that SOAT inhibition prevented crystal formation in cholesterol loaded RAW 264.7 macrophage cells. To our knowledge, this has not been reported before. While it is unclear what this implies, it is worthwhile to consider that other situations in which there is excess cholesterol may lead to SOAT1-dependent formation of cholesterol crystals. This may be relevant to subintimal macrophage and smooth muscle cells in patients with hypercholesterolemia or to neurons in patients with Niemann-Pick disease and other disorders affecting cholesterol metabolism. Moreover, it is possible that lipid droplet cholesterol crystals in hepatocytes affect subcellular interactions. In which case, defining the effect of such crystals, as well as the effect of preventing them from forming, on processes such as organelle organization and metabolism may provide more clarity on the role they play in MASH.

## Abbreviations

AAV: adeno-associated virus
ACAT: acyl co-A cholesterol acyltransferase
AcLDL: acetylated low-density lipoprotein
ANOVA: analysis of variance
BCA: bicinchoninic acid
Cas9: CRISPR-associated 9
CRE: cre recombinase
CRISPR: clustered regularly interspaced short palindromic repeats
DMEM: Dulbecco’s modified eagle medium
DMSO: dimethyl sulfoxide
ER: endoplasmic reticulum
GFP: green fluorescent protein
HepG2: human hepatoma G2
Hep3B: human hepatoma 3B
HFFC: high fat, fructose, cholesterol
LC3: light chain 3
LDL: low-density lipoprotein
MASH: metabolic dysfunction-associated steatohepatitis
MßCD: methyl-ß-cyclodextrin
mTOR: mechanistic/mammalian target of rapamycin
Nrf1/*Nfe2l1*: nuclear factor erythroid 2 related factor-1
Nrf2/*Nfe2l2*: nuclear factor erythroid 2 related factor-2
PBS: phosphate buffered saline
qPCR: quantitative polymerase chain reaction
SEM: standard error of the mean
SOAT: sterol O-acyltransferase
TBST: tris-buffered saline containing tween 20
TEM: transmission electron microscopy

