## Supplemental Figures for "Sterol O-acyltransferase (SOAT/ACAT) activity is required to form cholesterol crystals in hepatocyte lipid droplets"

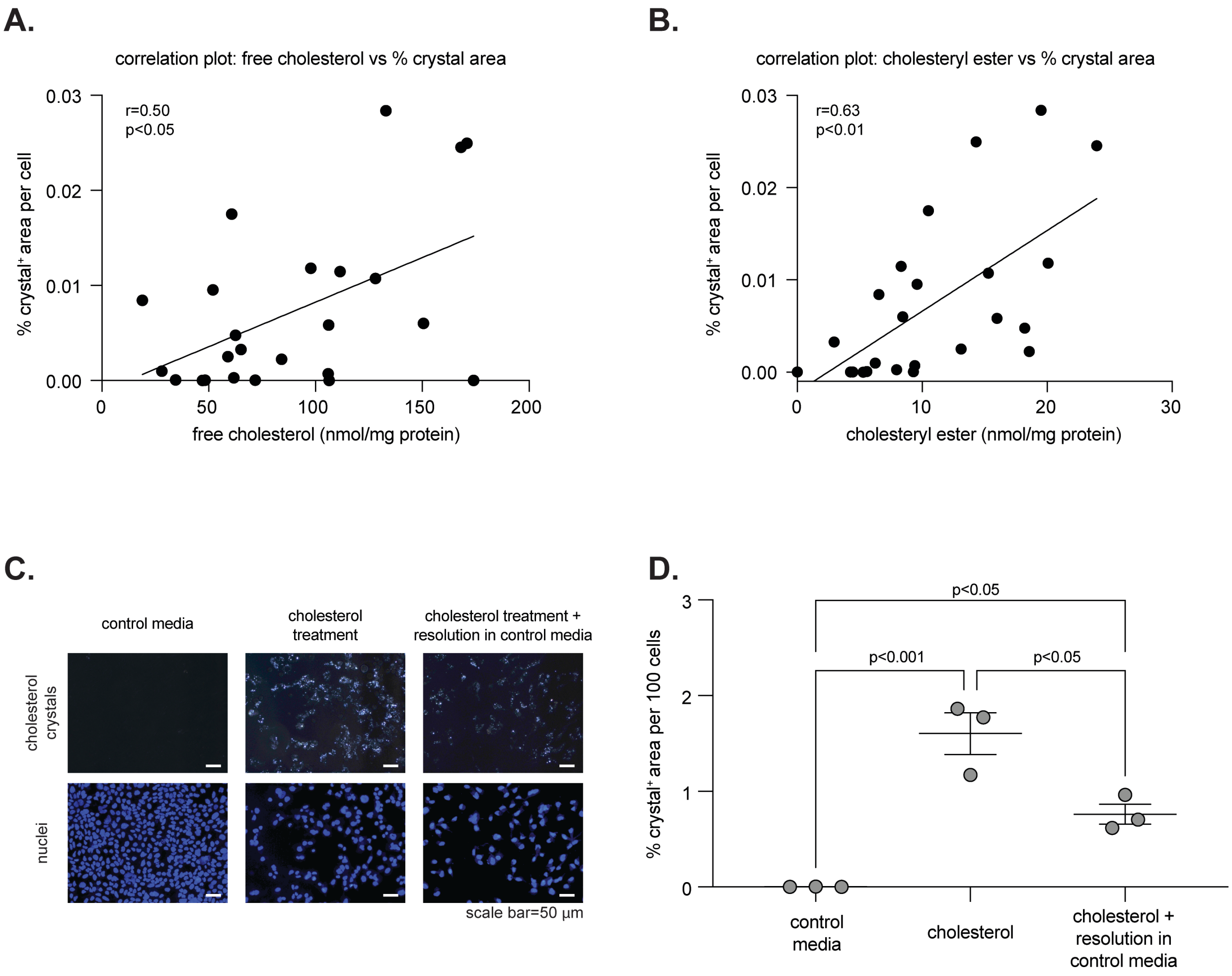


**Supplemental Figure 1. Validating the relationship between Hep3B cell cholesterol and crystal level.** (A and B) Correlation of crystals with free cholesterol (A) and cholesteryl ester (B). The individual data points and slope line are shown (n=24). The p-value and slope were determined using simple linear regression analysis. (C) Representative images of cells treated with control media, 200 µM cholesterol for 48 hours, or 200 µM cholesterol for 48 hours followed by control media for 48 hours. Detected crystals and nuclei is indicated in panel. (D) Quantified cholesterol crystals in (C) to examine whether formed crystals can resolve with respect to time (n=3). The p-value was determined by one-way analysis of variance, with Tukey's post-hoc test. Scale bar for (C) is shown in panel. Data in (D) are mean +/- standard error of the mean, with individual data points shown.

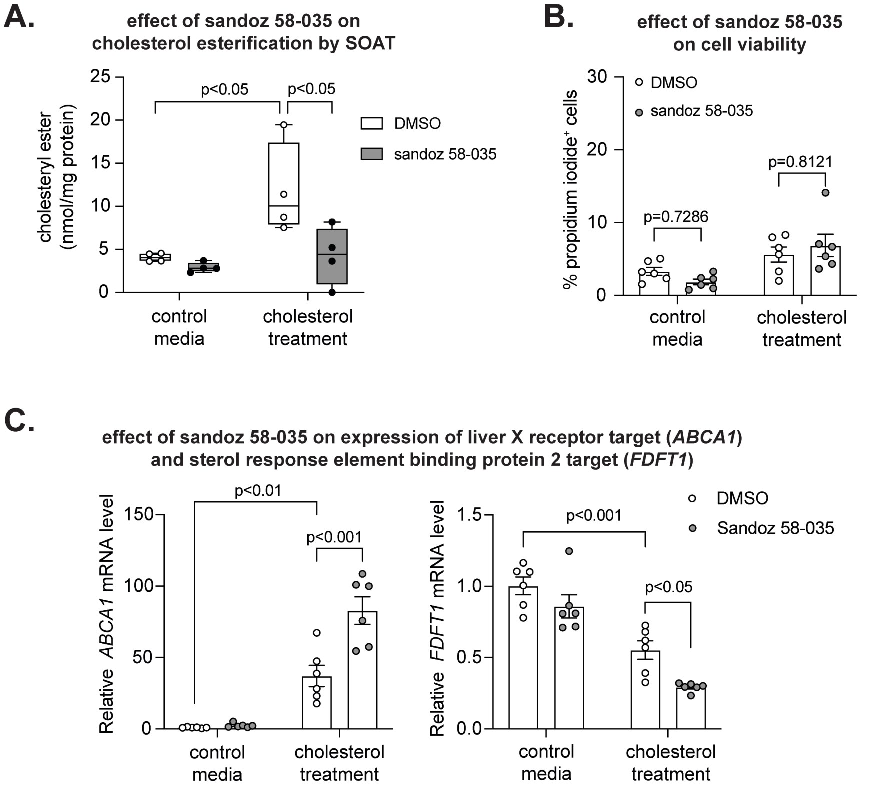


**Supplemental Figure 2. Effect of sandoz 58-035 on Hep3B cell cholesteryl ester, viability, and gene expression.** (A-C) Quantified levels of cholesteryl ester (A; n=3), dead cells (B; n=6), and expression of indicated cholesterol-regulated genes (C; n=6) in cells treated with or without cholesterol +/- sandoz 58-035. The p-value was determined by two-way analysis of variance, with Tukey's post-hoc test. Data in (A-C) are mean +/- standard error of the mean, with individual data points shown.

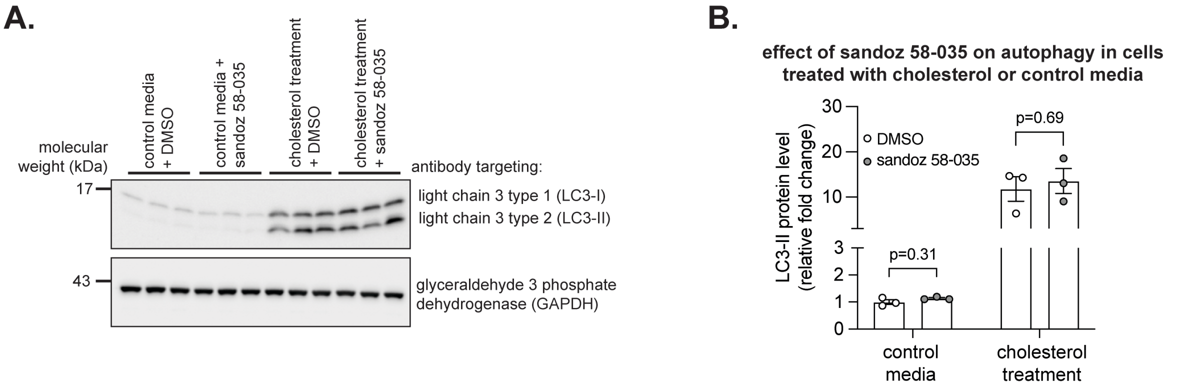


**Supplemental Figure 3. Effect of sandoz 58-035 on autophagy.** (A) Immunoblot of indicated proteins from Hep3B cells treated with or without cholesterol +/- sandoz 58-035. (B) Quantified immunoblot in (A) to examine the effect on autophagy (n=3). The p-value was determined by t-test, adjusted for multiple comparison. Data are mean +/- standard error of the mean, with individual data points shown.

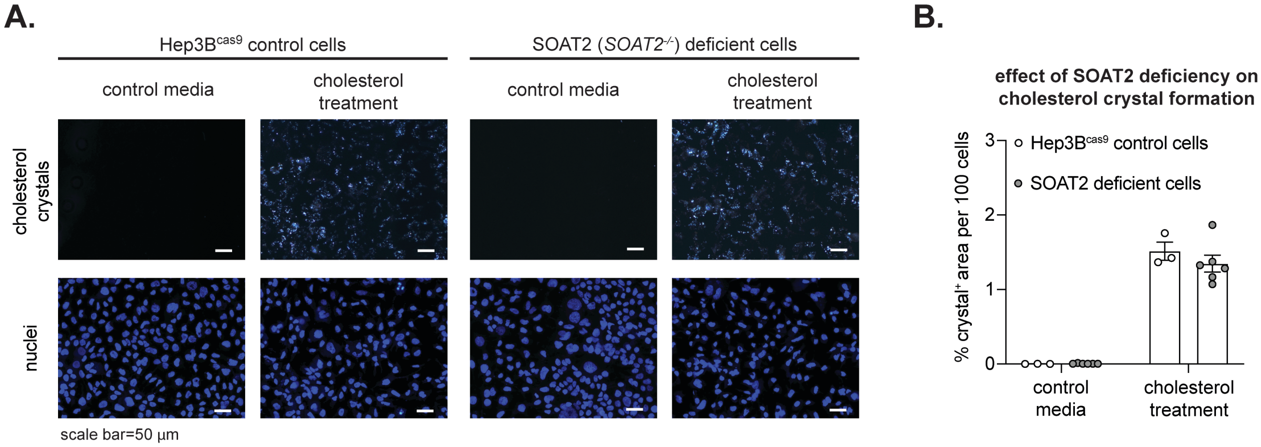


**Supplemental Figure 4. SOAT2 deficiency does not affect cholesterol crystal formation in Hep3B cells.** (A) Representative images of Hep3B$^{cas9}$ cells and SOAT2 deficient cells treated as indicated with or without cholesterol. (B) Quantified cholesterol crystals in (A) to examine the effect of SOAT2 deficiency on crystal formation (n=3-6). The p-value was determined by t-test, adjusted for multiple comparison. Scale bar for (A) is shown in panel. Data in (B) are mean +/- standard error of the mean, with individual data points shown.

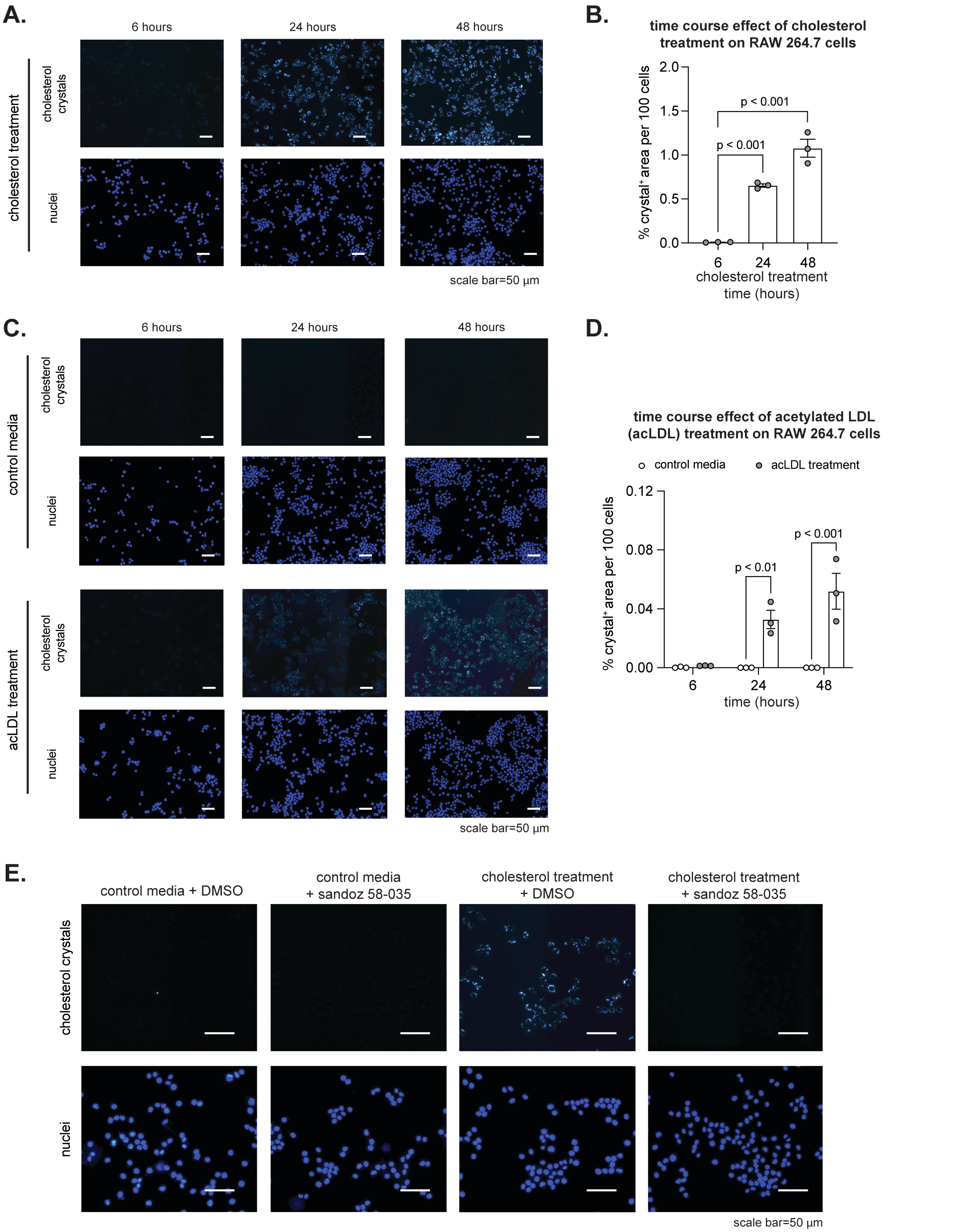


**Supplemental Figure 5. Time-dependent crystal formation in cholesterol loaded RAW 264.7 cells is prevented by sandoz 58-035.** (A and C) Representative images of RAW 264.7 cells treated with 200 µM cholesterol (A) or with control media or 50 µg/ml acetylated LDL (acLDL) (C) for 6-48 hours (n=3). (B and D) Quantified cholesterol crystals in (A) for (B) and (C) for (D) to examine effect of time on crystal formation (n=3). (E) Image of RAW 264.7 cells treated with or without 200 µM cholesterol +/- sandoz 58-035 to demonstrate that, like in Fig. 5C, sandoz 58-035 prevents crystals in cells loaded with cholesterol via MßCD-cholesterol complex method (n=1). The p-value in (B) was determined by one-way analysis of variance, with Dunnett's post-hoc test. The p-value in (D) was determined by t-test, adjusted for multiple comparison. Scale bar for (A), (C), and (E) is shown in panel. Data in (B) and (D) are mean +/- standard error of the mean, with individual data points shown.
